# Epidermal growth factor receptors in vascular endothelial cells contribute to functional hyperemia in the brain

**DOI:** 10.1101/2023.09.15.557981

**Authors:** Hannah R. Ferris, David C. Hill-Eubanks, Mark T. Nelson, George C. Wellman, Masayo Koide

## Abstract

Functional hyperemia – activity-dependent increases in local blood perfusion – underlies the on-demand delivery of blood to regions of enhanced neuronal activity, a process that is crucial for brain health. Importantly, functional hyperemia deficits have been linked to multiple dementia risk factors, including aging, chronic hypertension, and cerebral small vessel disease (cSVD). We previously reported crippled functional hyperemia in a mouse model of genetic cSVD that was likely caused by depletion of phosphatidylinositol 4,5-bisphosphate (PIP_2_) in capillary endothelial cells (EC) downstream of impaired epidermal growth factor receptor (EGFR) signaling. Here, using EC-specific EGFR-knockout (KO) mice, we directly examined the role of endothelial EGFR signaling in functional hyperemia, assessed by measuring increases in cerebral blood flow in response to contralateral whisker stimulation using laser Doppler flowmetry. Molecular characterizations showed that EGFR expression was dramatically decreased in freshly isolated capillaries from EC-EGFR-KO mice, as expected. Notably, whisker stimulation-induced functional hyperemia was significantly impaired in these mice, an effect that was rescued by exogenous administration of PIP_2_, but not by the EGFR ligand, HB-EGF. These data suggest that the deletion of the EGFR specifically in ECs depletes PIP_2_ and attenuates functional hyperemia, underscoring the central role of the endothelial EGFR signaling in cerebral blood flow regulation.

## 1 Introduction

Blood flow within the brain alters dynamically in response to spatial and temporal changes in neuronal activity, redirecting the delivery of blood-borne nutrients so as to support the elevated metabolic demands of active neurons. This phenomenon, termed functional hyperemia, is essential for maintaining brain health and computational activities, including cognition [1-4]. Functional hyperemia is mediated by an ensemble of neurovascular coupling mechanisms that translate increases in neuronal activity to increases in local blood flow through the integrated activity of multiple cell types of the neurovascular unit, comprising (in addition to neurons) astrocytes, vascular endothelial cells (ECs), vascular smooth muscle cells (SMCs), and pericytes [2, 5-10]. We recently reported that, among these cell types, brain capillary ECs, which are the building blocks of the vast, anastomosing network of capillaries – the smallest and most abundant vessels in the brain parenchyma – play a key role in linking neuronal activity to upstream arteriolar dilation [8]. They accomplish this by sensing focal neuronal activity in the form of modest increases in perivascular potassium ion (K^+^) concentrations caused by K^+^ efflux during each neuronal action potential. Increased levels of perivascular K^+^, in turn, activate capillary EC inward-rectifier Kir2.1 channels, which cause EC membrane potential hyperpolarization. Because Kir2.1 channels are also activated by membrane potential hyperpolarization, Kir2.1 channels in adjacent capillary ECs are also activated, resulting in propagation of the original hyperpolarizing (i.e., electrical) signal from EC to EC within the capillary bed. These signals ultimately reach upstream arterioles and arterioles, where they hyperpolarize overlying SMCs, causing arteriolar/artery dilation through a decrease in the open-state probability of voltage-dependent Ca^2+^ channels – a primary source of Ca^2+^ influx in vascular SMCs. This upstream arteriolar dilation results in increased local blood flow to the point of signal origin (i.e., region of heightened neuronal activity). Thus, capillary EC Kir2.1 channel activation is a keystone of neuronally triggered vasodilation, which is crucial for functional hyperemia in the brain [8].

In addition to being regulated by membrane potential and extracellular K^+^ concentration, Kir2.1 channel activity is also modulated by the plasma membrane phospholipid, phosphatidylinositol 4,5-bisphosphate (PIP_2_) [11, 12]. PIP_2_ is a requisite co-factor for Kir2.1 activity; thus, under PIP_2_-deficient conditions, such as in the context of increased PIP_2_ hydrolysis by Gq-type G-protein–coupled receptor (GqPCR) signaling-induced phospholipase C (PLC) activation, Kir2.1 channel activity is decreased [13]. Consistent with the idea that Kir2.1 channel activation is a critical step in capillary-initiated vasodilation, the attendant PIP_2_ depletion caused by activation of PLC through systemic administration of the GqPCR agonist, carbachol, decreases Kir2.1 currents in capillary ECs and attenuates capillary Kir2.1-mediated capillary-to-arteriole vasodilatory responses ex vivo and in vivo [13]. Notably, a PIP_2_ deficiency in capillary ECs has been shown to contribute to functional hyperemia deficits in a mouse model of CADASIL (cerebral autosomal dominant arteriopathy with subcortical infarcts and leukoencephalopathy) – the most common monogenic inherited form of cerebral small vessel diseases (cSVD) [14]. Strikingly, exogenous administration of PIP_2_ in CADASIL model mice rescues these deficits in capillary Kir2.1 channel currents, capillary-initiated arteriolar dilation, and functional hyperemia [14].

CADASIL is caused by mutations in the extracellular domain of the *NOTCH3* gene, expressed only in smooth muscle cells and pericytes in the brain [15, 16]. Previous studies have demonstrated that vascular pathologies in CADASIL involve accumulation of the matrix metalloproteinase inhibitor, TIMP3, in NOTCH3 extracellular domain (Notch3^ECD^) deposits located within the perivascular space surrounding brain vascular SMCs and capillary EC-wrapping pericytes. Suppression of the activity of the matrix metalloproteinase, ADAM17, by TIMP3 decreases shedding of the epidermal growth factor receptor (EGFR) ligand, heparin-binding EGF-like growth factor (HB-EGF), thereby decreasing EGFR activity [14, 17-19]. Interestingly, application of exogenous HB-EGF rescues capillary EC Kir2.1 currents, capillary-to-arteriole vasodilatory signaling, and functional hyperemia in CADASIL mice – effects comparable to those produced by PIP_2_ administration [14]. These findings suggest that down-regulated capillary EC EGFR signaling promotes a reduction in plasma membrane PIP_2_ levels, in turn decreasing Kir2.1 channel activity, crippling capillary-initiated vasodilatory responses, and impairing functional hyperemia.

Here, using a newly developed EC-EGFR-knockout (KO) mice, we sought to elucidate the role of endothelial EGFR signaling in functional hyperemia. To this end, we evaluated functional hyperemia, measured as an increase in cerebral blood flow (CBF) in the barrel cortex in response to contralateral whisker stimulation, using laser Doppler flowmetry. We further tested whether treatment with the Kir2.1 channel co-factor PIP_2_ or the EGFR ligand HB-EGF altered functional hyperemia in EC-EGFR-KO mice.

## 2. Results

### 2.1 Characterization of capillary ECs from EC-EGFR-KO mice

To characterize the newly developed inducible EC-EGFR-KO mouse model, we first confirmed ablation of EGFR protein in brain capillary ECs of these mice. Capillaries were isolated and collected from the brain cortex using a modification of the previously described protocol [20], as detailed in Methods. The purity of isolated capillary samples was verified using a dual-promoter double-color (acta2-RCaMP/cdh5-GCaMP8) mouse strain, which expresses the red fluorescent protein (RCaMP) in SMCs and pericytes, and the green fluorescent protein (GCaMP8) in ECs. As shown in Figure 1A, the vast majority (>98%) of microvessels obtained from these EC/SMC dual-reporter mice using this isolation protocol were GCaMP-positive and RCaMP-negative, demonstrating that these samples contained capillary ECs and were largely devoid of SMCs. Next, using freshly isolated capillary ECs samples from EC-EGFR-KO and WT mice, we quantified EGFR protein using a sandwich enzyme-linked immunosorbent assay (ELISA). Total EGFR protein expression in capillary ECs was dramatically (∼5-fold) decreased in EC-EGFR-KO mice (4.2 ± 0.7 ng/mg; n = 6) compared with that in wild-type (WT), Cre-negative littermates (22.1 ± 2.7 ng/mg; n=8) (Figure 1B). No differences in body weight, blood pressure, or heart rate were observed between EC-EGFR-KO and WT animals (Table 1), consistent with previously reported data from a strain of EC-EGFR-KO mice generated using the non-inducible Tie2-promoter [21].

**Table 1:** Physiological parameters in EC-EGFR-KO mice.

|  | WT (n=8) | EC-EGFR-KO (n=8) |
| --- | --- | --- |
| Age (weeks old) | 20.2 ± 2.2 | 20.8 ± 1.3 |
| Body weight (g) | 26.8 ± 1.2 | 27.3 ± 1.4 |
| Blood pressure (mmHg) | 91.3 ± 2.9 | 91.8 ± 1.1 |
| Heart rate (bpm) | 498 ± 8 | 491 ± 6 |

**Figure 1.**
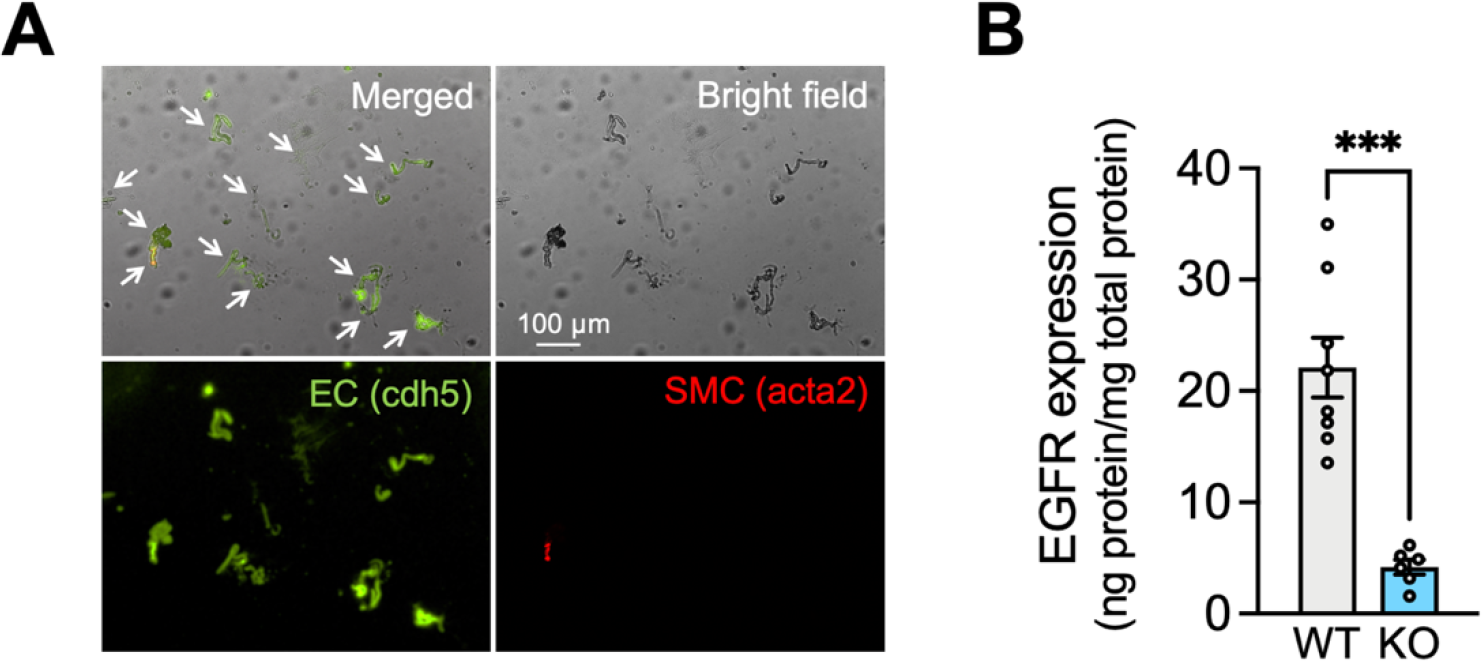
Characterization of capillary ECs from EC-EGFR-KO mice. **A:** Bright field and fluorescent images of isolated microvessels from EC/SMC dual-reporter mice. Green fluorescent protein expresses in ECs under the cadherin-5 (cdh5) promotor, and red fluorescent protein expresses in SMC/pericytes under the acta2 promoter. The vast majority (>98%) of tissue obtained was green fluorescent-positive and red fluorescent-negative microvessels, demonstrating that the collected tissue samples were capillaries. **B:** EGFR protein in freshly isolated capillaries, pooled from 4 animals for each sample, were quantified by ELISA. EGFR protein was dramatically decreased in EC-EGFR-KO mice compared to WT animals (i.e., Cre-negative littermates). Data are presented as mean ± SEM (n = 8 in WT, n = 6 in KO). *** P < 0.001 between groups by unpaired t-test.

### 2.2 Impaired functional hyperemia in EC-EGFR-KO mice

We next assessed functional hyperemia in EC-EGFR-KO and WT mice using laser-Doppler flowmetry. As shown in Figure 2 (A and B), contralateral whisker stimulation caused a 29.2% ± 1.1% increase in CBF in WT animals (n = 8); by comparison, the same stimulation caused a significantly attenuated (∼50%) hyperemic response (15.4 ± 0.7% increase) in EC-EGFR-KO mice (n = 8). To evaluate the contribution of Kir2.1 channel-mediated signaling, we topically applied the selective Kir2.1 channel blocker, Ba^2+^ (100 μM), to the superfusate flowing over the somatosensory cortex [8, 14, 22]. Consistent with the previously established involvement of Kir2.1 channels in the functional hyperemic response to whisker stimulation [8, 14, 22], this treatment substantially decreased functional hyperemia in both EC-EGFR-KO and WT mice, leaving the same residual response in both genotypes (Figure 2C). However, the Kir2.1-mediated (i.e., Ba^2+^-sensitive) component of this reduction was greater in WT mice (70.9% ± 2.5% reduction; n = 8) than in EC-EGFR-KO mice (46.8% ± 2.9% reduction; n = 8) (Figure 2D), suggesting that EC-specific EGFR ablation eliminated much of the Kir2.1-dependent response. Collectively, these observations indicate that ablation of EC-EGFR impairs Kir2.1-mediated capillary-to-arteriole vasodilatory signaling.

**Figure 2:**
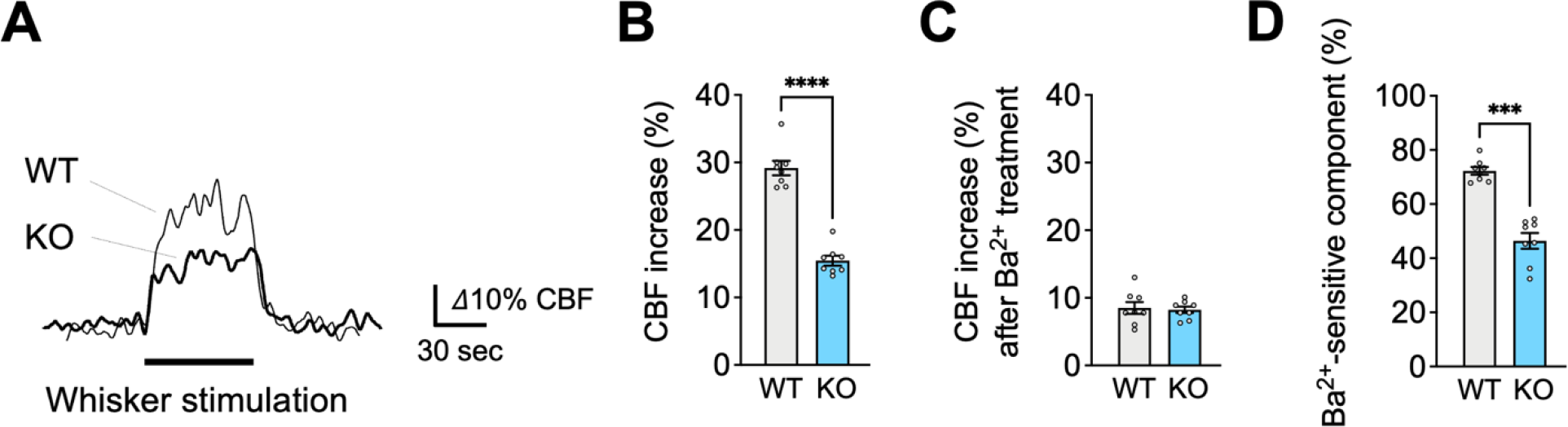
Impaired functional hyperemia in EC-EGFR-KO mice. **A:** Representative traces showing CBF increases in contralateral somatosensory cortex during whisker stimulation. Cre-negative littermates (WT) were used as a control group in comparison to EC-EGFR-KO (KO) mice. **B:** Summary data of whisker stimulation-induced functional hyperemia. **C:** CBF increase during whisker stimulation after treatment with Ba^2+^ (100 μM), a Kir2.1 channel blocker. **D:** Ba^2+^ sensitive-component of functional hyperemia, i.e., the difference in responses before and after Ba^2+^ treatment, indicating the contribution of the Kir2.1 channels to whisker stimulation-induced functional hyperemia. Data are presented as mean ± SEM (n = 8 in WT, n = 8 in KO). **** P < 0.0001, *** P < 0.001 between groups by unpaired t-test.

### 2.3 PIP_2_, but not HB-EGF, restores functional hyperemia in EC-EGFR-KO mice

To further explore the basis of Kir2.1 channel impairment in EC-EGFR-KO mice, we examined the ability of PIP_2_, an endogenous Kir2.1 co-factor, to reverse functional hyperemia deficits in these mice. To this end, we administered the synthetic PIP_2_ analog, dipalmitoyl-PIP_2_ (0.5 mg/kg body weight), to EC-EGFR-KO mice or WT mice through a catheter placed in the femoral artery [14] and tested whisker stimulation-induced functional hyperemia 20 minutes later. As shown in Figure 3 (A and B), this maneuver significantly increased functional hyperemic responses in EC-EGFR-KO mice (23.6% ± 1.6% increase in CBF; n = 4) compared with those elicited prior to PIP_2_ administration (15.3% ± 0.7% increase in CBF). Consistent with previous observations [14], PIP_2_ administration did not impact functional hyperemic responses to whisker stimulation in WT animals (28.1% ± 1.0% and 27.2 ± 1.8% before and after PIP_2_, respectively; n = 4) (Figure 3B). Interestingly, PIP_2_ administration enhanced the Kir2.1-mediated (Ba^2+^-sensitive) component in EC-EGFR-KO mice, increasing it to a level comparable to that observed in WT mice (Figure 3C), confirming that PIP_2_ supplementation improved Kir2.1 channel function. We next tested whether treatment with the EGFR ligand, HB-EGF, which rescues functional hyperemic responses in CADASIL model mice [18, 19], could similarly normalize functional hyperemia in EC-EGFR-KO mice. Consistent with the absence of its cognate receptor, HB-EGF (30 ng/mL), topically applied for 20 minutes [14, 18], did not alter functional hyperemic responses in EC-EGFR-KO mice, which exhibited persistent functional hyperemia deficits despite HB-EGF cortical superfusion (with HB-EGF, 14.0% ± 1.6%; without HB-EGF, 16.0% ± 0.8%; n = 4). The Ba^2+^-sensitive component also remained significantly smaller in EC-EGFR-KO mice (Figure 4C). Functional hyperemia was also unaffected by HB-EGF treatment in WT animals, which presented similar increases in cortical blood flow in response to whisker stimulation before and after HB-EGF (with HB-EGF, 27.7% ± 2.6%; without HB-EGF, 26.7% ± 1.5%; n = 4). These data suggest that PIP_2_ depletion contributes to the decreased activity of capillary EC Kir2.1 channels and functional hyperemia deficits in EC-EGFR-KO mice.

**Figure 3:**
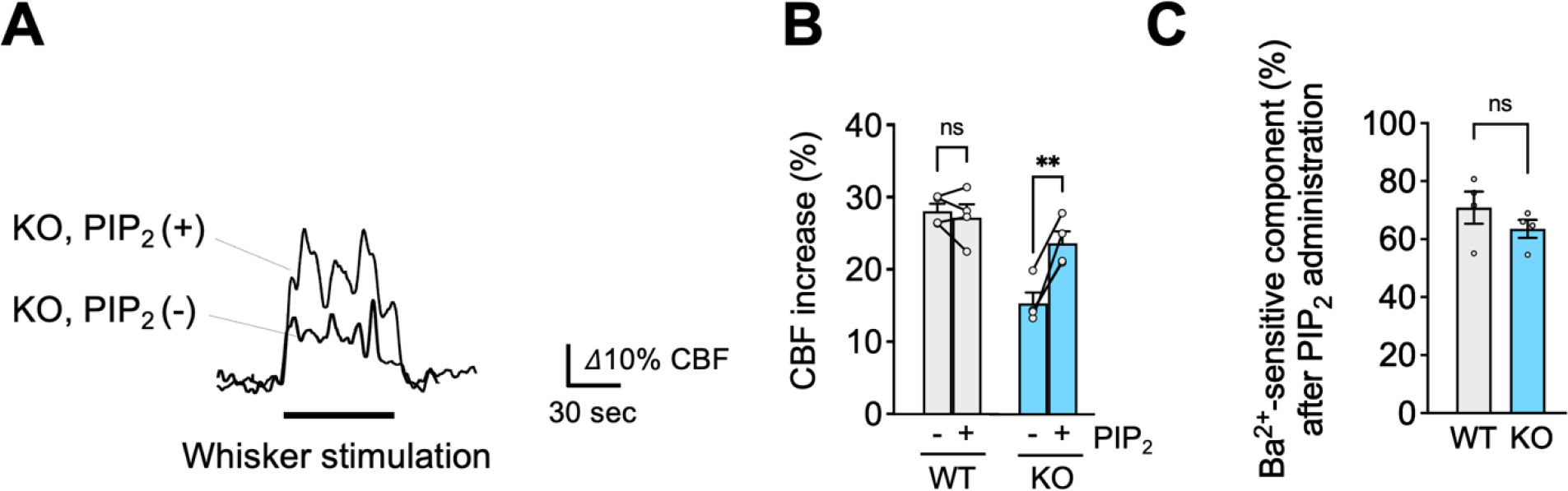
PIP_2_, an endogenous Kir2.1 channel co-factor, restored functional hyperemia in EC-EGFR-KO mice. **A:** Whisker stimulation-induced functional hyperemia before and after PIP_2_ treatment in EC-EGFR-KO mice. **B:** Summary data showing PIP_2_ treatment restores functional hyperemia deficits in EC-EGFR-KO mice, with little impact on WT animals. **C:** Ba^2+^ sensitive-component of functional hyperemia after PIP_2_ treatment. Data are presented as mean ± SEM (n = 4 in WT, n = 4 in KO). ** P < 0.01, ns: not significant between groups by paired t-test (B) or by unpaired t-test (C).

**Figure 4:**
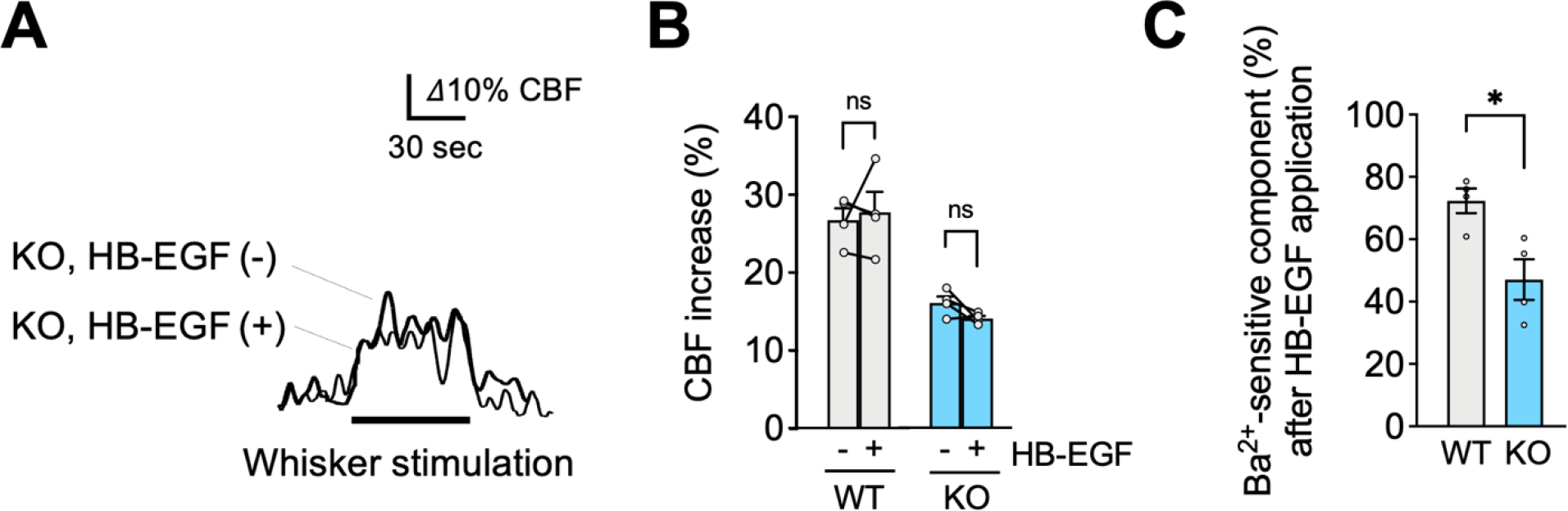
HB-EGF, an EGFR ligand, failed to restore functional hyperemia in EC-EGFR-KO mice. **A:** Whisker stimulation-induced functional hyperemia before and after HB-EGF treatment in EC-EGFR-KO mice. **B:** Summary data showing that HB-EGF treatment did not alter whisker stimulation-induced functional hyperemia in either EC-EGFR-KO or WT mice. **C:** Ba^2+^ sensitive-component of functional hyperemia after HB-EGF treatment. Data are presented as mean ± SEM (n = 4 in WT, n = 4 in KO). * P < 0.05, ns: not significant between groups by paired t-test (B) or by unpaired t-test (C).

**Figure 5:**
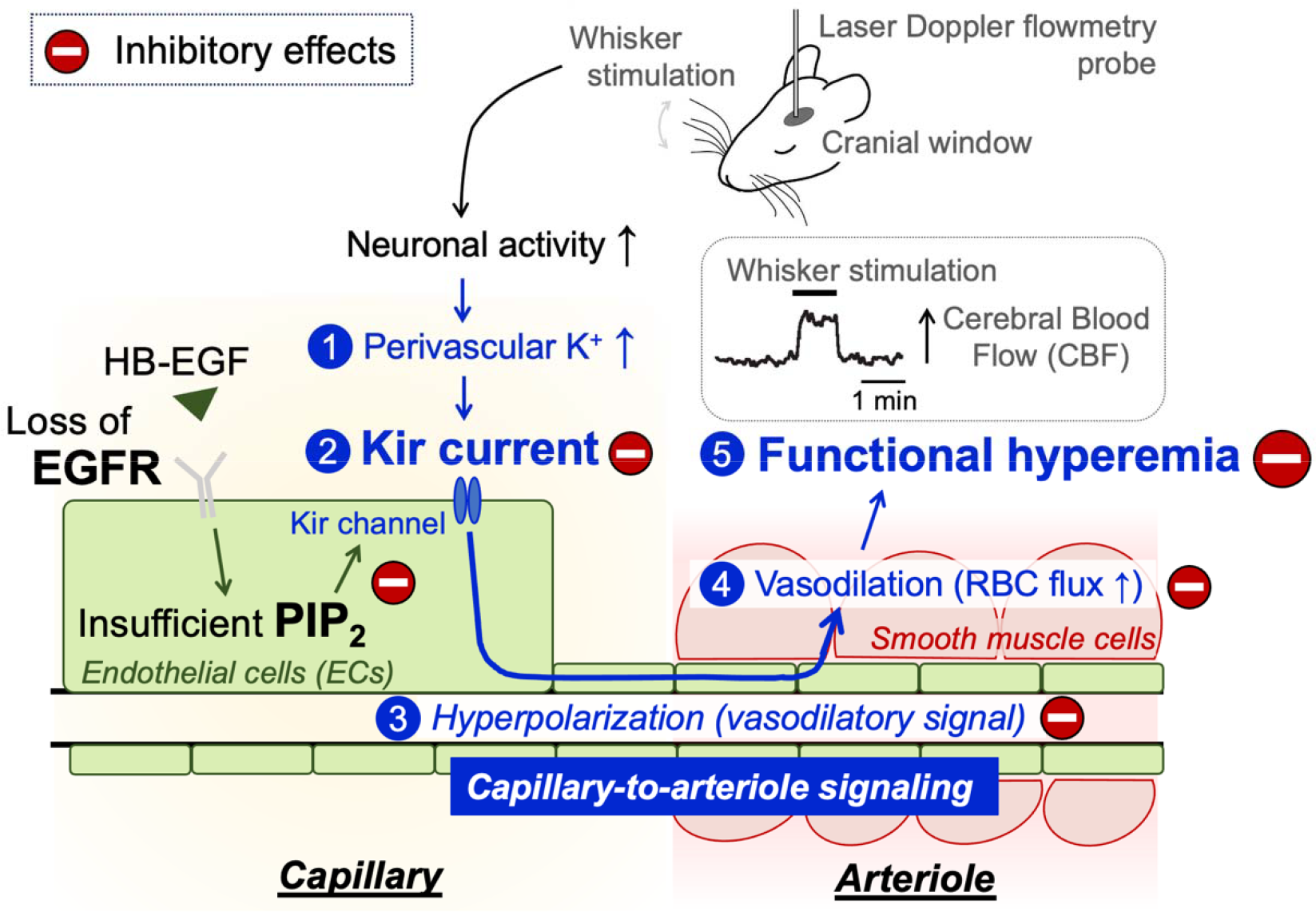
Schematic illustration of the proposed signaling pathway underlying functional hyperemia deficits in EC-EGFR-KO mice. Whisker stimulation and subsequent neuronal activation, which increases perivascular K^+^ during neuronal repolarization, results in Kir2.1 channel activation in capillary ECs. Kir2.1 channel-initiated hyperpolarizing vasodilatory signal rapidly propagates upstream and dilates the precapillary arteriole, increasing downstream tissue perfusion. This local increase in blood flow in neuronally active regions of the brain is termed functional hyperemia, which is detected using laser Doppler flowmetry in this study. Genetic deletion of EGFR disrupts functional hyperemia in response to whisker stimulation by causing Kir2.1 channel dysfunction, which is likely attributed to lowering the level of PIP_2_, an essential co-factor of Kir2.1 channels.

## 3. Discussion

In this study, we examined the role of endothelial EGFR signaling in functional hyperemic responses in the brain using newly-developed EC-specific EGFR-KO mice. EGFR, originally identified as a cancer-promoting protein [23], is now known to be involved in a multitude of (patho)physiological phenomena [24-27], including functional hyperemia in the brain [14, 18]. Here, we validated the loss of EGFR protein in capillaries of EC-EGFR-KO mice and show that the selective ablation of endothelial EGFR disrupts capillary-to-arteriole signaling and significantly impairs functional hyperemia. These findings are consistent with our prior work using CADASIL model mice harboring mutations in the NOTCH3 gene, which indicated that functional hyperemia is attenuated by EGFR-dependent down-regulation of Kir2.1 activity in ECs [14]. Importantly, in CADASIL mice, Kir2.1 channel dysfunction is observed only in capillary ECs, and not in arteriolar SMCs [14]. Moreover, functional hyperemia deficits in CADASIL mice were rescued by treatment with HB-EGF [14], an observation that suggested that disabled endothelial EGFR signaling is a major pathology in functional hyperemia deficits in CADASIL. Collective with the current study, these data confirm that endothelial EGFR signaling is an important contributor to functional hyperemia in the brain.

Functional hyperemia deficits in EC-EGFR-KO mice are likely attributable to crippled Kir2.1 channel-mediated signaling, as supported by the diminished Ba^2+^ sensitive-component of whisker-stimulated increases in CBF in these mice (Figure 2D). This is consistent with a previous report that loss of Kir2.1 channel function (e.g., through genetic deletion of Kir2.1 channels in ECs [8]) leads to functional hyperemia deficits. In CADASIL model mice, Kir2.1 dysfunction was shown to result from an insufficiency of the Kir2.1 channel co-factor, PIP_2_ [14]; this study further showed that treatment with exogenous PIP_2_ substantially restored Kir2.1 channel currents in capillary ECs and rescued functional hyperemia deficits [14]. Similar to the case for CADASIL mice, functional hyperemia deficits in EC-EGFR-KO mice were alleviated by the administration of PIP_2_ (Figure 3). These findings suggest that elimination of endothelial EGFRs reduces PIP_2_ levels, subsequently impairing functional hyperemia by diminishing Kir2.1 channel activity. Further, the observations that exogenous PIP_2_ or HB-EGF did not cause additional increases in functional hyperemia response indicate that EGFR-intact animals hold sufficient PIP_2_ in their capillary EC plasma membrane to sustain Kir 2.1 channel functionality. The mechanism by which the EGFR maintains PIP_2_ levels remains to be elucidated, although it has been reported that EGFR activation enhances the production of ATP [28, 29], which is needed at relatively high (generally micromolar) concentrations to support the synthesis of PIP_2_ [30]. Thus, elimination of EGFRs in ECs may dampen ATP production and levels, and diminish PIP_2_ synthesis, resulting in PIP_2_-depleted conditions and decreased Kir2.1 channel activity.

Our results indicate that the endothelial EGFR signaling is a requisite component of functional hyperemia in the brain, likely through the maintenance of capillary Kir2.1 channel function. However, this work does not exclude a contribution of smooth muscle EGFR signaling in neuronal activity-dependent increases in local blood perfusion. In cerebral arteries [31] and arterioles [19], activation of smooth muscle EGFR by ligands, such as HB-EGF, promotes vasoconstriction via suppression of voltage-gated potassium (K_V_1.5) channel currents, leading to membrane depolarization and increased intracellular Ca^2+^ [19, 32-34]. Thus, activation of smooth muscle EGFRs causes arterial constriction directly [19, 35] and/or enhances agonist-induced contractile responses [36]. In CADASIL, which results from mutant Notch3^ECD^ accumulation-dependent dysregulation of TIMP3/ADAM17/HB-EGF/EGFR signaling, the decreased EGFR signaling is not limited to ECs but also impacts arterial SMCs. Hence, intraluminal pressure-induced cerebral artery constriction (myogenic tone), a mechanism intrinsic to SMCs, is greatly attenuated in CADASIL mice, which can be recovered by treatment with HB-EGF [19]. Notably, functional hyperemia deficits in CADASIL mice were restored by the EGFR ligand, HB-EGF. Therefore, it is conceivable that HB-EGF restores functional hyperemic responses in CADASIL mice by improving both endothelial EGFR signaling (through enhanced capillary Kir2.1 channel function), as well as smooth muscle EGFR signaling (by impacting arterial contractility and/or vasodilatory capability). Nonetheless, this present study clearly demonstrates that selective loss of EC EGFR signaling is sufficient to disrupt functional hyperemia, consistent with endothelial EGFR signaling playing a key role in the regulation of focal vasodilatory responses within active regions of the brain.

## 4. Material and Methods

### Animals

<u>EC-EGFR-KO mice</u> were obtained by crossbreeding cdh5-CreERT2 mice (obtained from Dr. Ralf Adams, Cancer Research UK London Research Institute) [37] and floxed-EGFR (kindly gifted from Dr. David Thredgill, Texas A&M University) [38]. Cadherin-5 (cdh5, aka VE-cadherin) is a vascular endothelial cell-specific adhesion molecule and was employed as an EC-specific promoter [37]. Cdh5-CreERT2 -positive, floxed-EGFR homozygous offspring, including both males and females, were treated with tamoxifen by adding it to their chow (Envigo, #TD.130859, ∼40 mg/kg body weight/day) at 8 weeks of age for 7 days. Animals were used experimentally 2-4 weeks after ending tamoxifen treatment to allow for extinction of pre-existing EGFR protein. Cre-negative littermates were used as control “wildtype (WT)” animals. <u>EC/SMC dual-reporter mice</u> were obtained by crossbreeding of acta2-RCaMP (JAX, strain #028345; CHROMus) [39] and cdh5-GCaMP8 (JAX, strain #33342; CHROMus) [40]. This dual-promoter double-color mouse strain has red fluorescent protein in acta2-positive cells (i.e., smooth muscle cells and pericytes [9]) and green fluorescent protein in cdh5-positive cells (i.e., endothelial cells) and was used to distinguish these cell types [40].

### Isolation of brain parenchymal capillaries

Capillaries were collected from both cerebral cortices using a modified protocol that was previously described for collection of cerebral microvessels [20]. Briefly, pia membrane and brain surface vessels were physically removed from isolated cortices using a Kimwipe. Cortices were then homogenized in ice-cold artificial cerebrospinal fluid (aCSF: 125 mM NaCl, 3 mM KCl, 26 mM NaHCO_3_, 1.25 mM NaH_2_PO_4_, 1 mM MgCl_2_, 4 mM glucose, 2 mM CaCl_2_, pH 7.3) using a Dounce homogenizer and homogenates centrifuged at 2,000 x*g* for 5 min at 4°C. Crude tissue precipitates were re-suspended in 17.5% dextran (∼ 70 kDa, in aCSF) and centrifuged at 4,000 *x g* for 20 min at 4°C, which allowed separation of a blood vessel precipitate from an upper layer of myelin. Next, capillaries were isolated from blood vessel precipitate by a two-step filtration process. First, the vasculature-rich fraction was re-suspended in aCSF and filtered through a 200-μm mesh strainer, removing large vessels that were retained in the filter. The remaining suspension was then filtered through a 40-μm mesh strainer. Capillary-rich microvessels retained on the 40-μm mesh strainer were retrieved by washing off the inverted strainer with aCSF, and collected by centrifuging at 2,000 x *g* for 5 min. The vast majority (>98%) of tissue obtained by this method were capillaries, which lack smooth muscle cells (Fig. 1A).

### Measurement of EGFR expression

EGFR protein was quantified in cortical capillary preparations by ELISA (Invitrogen, EM23RB) following the manufacturer’s instruction. Freshly isolated and collected capillaries, pooled from 4 animals for each sample, were lysed in 200 μL of T-PER™ tissue protein extraction reagent (ThermoFisher) containing protease inhibitors (1 mM AEBSF, 0.8 μM aprotinin, 50 μM Bestatin, 15 μM E-64, 5 mM EDTA, 20 μM leupeptin, 10 μM pepstatin A). Samples (90 μL per well, in duplicates) or standard protein (mouse recombinant EGFR, provided in the assay kit) were applied to anti-mouse EGFR antibody-coated 96-well plates and incubated overnight at 4°C. After washing the plate with kit-provided wash buffer, biotin conjugated anti-mouse EGFR antibody was added to the plate and incubated for 1 hr at room temperature (RT) with gentle shaking. The plate was then rinsed with wash buffer, treated with horseradish peroxidase-conjugated streptavidin (for 45 min at RT), washed again, and 3,3’,5,5’-tetramethylbenzidine, a peroxidase substrate, was applied for 30 min at RT in the dark. Peroxidase reactions were terminated by adding the Stop solution, and optical absorbance at 450 nm was immediately measured by a 96-well plate reader (Enzo). EGFR protein expression (average of duplicate measurements) was normalized by total protein concentration, determined by Bradford assay for each sample.

### Laser Doppler flowmetry

Functional hyperemia was measured by laser Doppler flowmetry as increases in CBF in response to contralateral whisker stimulation as described previously [8, 14, 22]. Briefly, under a surgical plane of isoflurane-anesthesia, a catheter was inserted in a femoral artery for blood pressure monitoring and a cranial window was prepared over the somatosensory cortex. Cranial windows were superfused with aCSF (aerated with 5% CO_2_/20% O_2_/75% N_2_, warmed to 37°C). After switching anesthesia to a combination of urethane (750 mg/kg, i.p.) and alpha chloralose (50 mg/kg, i.p.), CBF was monitored via a laser Doppler flow probe (PeriMed) placed over the somatosensory cortex through the cranial window. Functional hyperemia was induced by stroking the opposite side vibrissae at a frequency of 3-5 Hz for 1 minute (whisker stimulation) and expressed as a percent increase in CBF. Drugs were applied topically to the somatosensory cortex via aCSF superfusate to the cranial window, except when specifically stated. Throughout CBF measurements, blood pressure and heart rate were monitored through a femoral arterial cannula, and body temperature was maintained at 37°C using a heating pad thermostatically controlled by a rectal probe. All data were recorded and analyzed using LabChart software (AD Instruments).

## 5. Conclusion

This study used a newly developed EC-EGFR-KO mouse model to demonstrate that the endothelial EGFR contributes to functional hyperemia, an essential CBF regulatory mechanism. In contrast to EGFRs in vascular SMCs, which play a role in arterial contractility, endothelial EGFR signaling modulates focal vasodilatory responses in neuronally active regions of the brain. Disrupting endothelial EGFR signaling impairs functional hyperemia, likely by incapacitating Kir2.1 channel functionality in capillaries. Overall, our study contributes to a deeper understanding of the mechanisms responsible for functional hyperemia deficits in cSVD/CADASIL model mice. Moving forward, investigations into vascular EGFR signaling, involving both SMCs and ECs, promise to shed additional light on the pathologies associated with cerebral vascular dysfunction occurring in a wide variety of diseases.

## Author contributions

MK designed and directed the study, evaluated functional hyperemia, and prepared the manuscript, HF managed EC-EGFR-KO mouse development and measured EGFR expression, MTN obtained and provided cdh5-CreERT2 mice, and GCW and DHE edited the manuscript. All authors approved the manuscript submission.

## Funding

This work was supported by the Orphan Disease Center Million Dollar Bike Ride Pilot Grant Program (MDBR-21-101-CADASIL to MK), the American Heart Association (14SDG20150027 to MK), and National Institute of Health (P20-GM-135007 to MK and MTN; R35-HL-140027, R01-NS-110656, and RF1-NS-128963 to MTN, and R01-HL-142888 to GCW).

## Acknowledgement

We would like to thank Drs. David Thredgill (Texas A&M University), Ralf Adams (Cancer Research UK London Research Institute, United Kingdom; University of Münster, Germany), and Mike Kotlikoff (Cornell University; CHROMus) for their generous sharing of transgenic mice for this study.

## Conflict of Interest

All authors have no conflict to declare.

## Notes

### Competing Interest Statement

The authors have declared no competing interest.

## References

[1] Iadecola C. The Neurovascular Unit Coming of Age: A Journey through Neurovascular Coupling in Health and Disease. Neuron. 2017;96:17–42.

[2] Schaeffer S, Iadecola C. Revisiting the neurovascular unit. Nat Neurosci. 2021.

[3] Lecrux C, Hamel E. The neurovascular unit in brain function and disease. Acta Physiol (Oxf). 2011;203:47–59.

[4] Santisteban MM, Iadecola C, Carnevale D. Hypertension, Neurovascular Dysfunction, and Cognitive Impairment. Hypertension. 2023;80:22–34.

[5] Haydon PG, Carmignoto G. Astrocyte control of synaptic transmission and neurovascular coupling. Physiol Rev. 2006;86:1009–31.

[6] Iadecola C, Nedergaard M. Glial regulation of the cerebral microvasculature. Nat Neurosci. 2007;10:1369–76.

[7] Filosa JA, Bonev AD, Straub SV, Meredith AL, Wilkerson MK, Aldrich RW, et al. Local potassium signaling couples neuronal activity to vasodilation in the brain. Nat Neurosci. 2006;9:1397–403.

[8] Longden TA, Dabertrand F, Koide M, Gonzales AL, Tykocki NR, Brayden JE, et al. Capillary K(+)-sensing initiates retrograde hyperpolarization to increase local cerebral blood flow. Nat Neurosci. 2017;20:717–26.

[9] Gonzales AL, Klug NR, Moshkforoush A, Lee JC, Lee FK, Shui B, et al. Contractile pericytes determine the direction of blood flow at capillary junctions. Proc Natl Acad Sci U S A. 2020;117:27022–33.

[10] Sancho M, Klug NR, Mughal A, Koide M, Huerta de la Cruz S, Heppner TJ, et al. Adenosine signaling activates ATP-sensitive K(+) channels in endothelial cells and pericytes in CNS capillaries. Sci Signal. 2022;15:eabl5405.

[11] Hansen SB, Tao X, MacKinnon R. Structural basis of PIP2 activation of the classical inward rectifier K+ channel Kir2.2. Nature. 2011;477:495–8.

[12] Huang CL, Feng S, Hilgemann DW. Direct activation of inward rectifier potassium channels by PIP2 and its stabilization by Gbetagamma. Nature. 1998;391:803–6.

[13] Harraz OF, Longden TA, Dabertrand F, Hill-Eubanks D, Nelson MT. Endothelial GqPCR activity controls capillary electrical signaling and brain blood flow through PIP2 depletion. Proc Natl Acad Sci U S A. 2018;115:E3569–E77.

[14] Dabertrand F, Harraz OF, Koide M, Longden TA, Rosehart AC, Hill-Eubanks DC, et al. PIP(2) corrects cerebral blood flow deficits in small vessel disease by rescuing capillary Kir2.1 activity. Proc Natl Acad Sci U S A. 2021;118.

[15] Chabriat H, Joutel A, Dichgans M, Tournier-Lasserve E, Bousser MG. Cadasil. Lancet Neurol. 2009;8:643–53.

[16] Joutel A, Corpechot C, Ducros A, Vahedi K, Chabriat H, Mouton P, et al. Notch3 mutations in CADASIL, a hereditary adult-onset condition causing stroke and dementia. Nature. 1996;383:707–10.

[17] Joutel A, Monet-Leprêtre M, Gosele C, Baron-Menguy C, Hammes A, Schmidt S, et al. Cerebrovascular dysfunction and microcirculation rarefaction precede white matter lesions in a mouse genetic model of cerebral ischemic small vessel disease. J Clin Invest. 2010;120:433–45.

[18] Capone C, Dabertrand F, Baron-Menguy C, Chalaris A, Ghezali L, Domenga-Denier V, et al. Mechanistic insights into a TIMP3-sensitive pathway constitutively engaged in the regulation of cerebral hemodynamics. Elife. 2016;5.

[19] Dabertrand F, Krøigaard C, Bonev AD, Cognat E, Dalsgaard T, Domenga-Denier V, et al. Potassium channelopathy-like defect underlies early-stage cerebrovascular dysfunction in a genetic model of small vessel disease. Proc Natl Acad Sci U S A. 2015;112:E796–805.

[20] Lee YK, Uchida H, Smith H, Ito A, Sanchez T. The isolation and molecular characterization of cerebral microvessels. Nat Protoc. 2019;14:3059–81.

[21] Schreier B, Stern C, Dubourg V, Nolze A, Rabe S, Mildenberger S, et al. Endothelial epidermal growth factor receptor is of minor importance for vascular and renal function and obesity-induced dysfunction in mice. Sci Rep. 2021;11:7269.

[22] Koide M, Harraz OF, Dabertrand F, Longden TA, Ferris HR, Wellman GC, et al. Differential restoration of functional hyperemia by antihypertensive drug classes in hypertension-related cerebral small vessel disease. J Clin Invest. 2021;131:e149029.

[23] Merlino GT, Xu YH, Ishii S, Clark AJ, Semba K, Toyoshima K, et al. Amplification and enhanced expression of the epidermal growth factor receptor gene in A431 human carcinoma cells. Science. 1984;224:417–9.

[24] Avraham R, Yarden Y. Feedback regulation of EGFR signalling: decision making by early and delayed loops. Nat Rev Mol Cell Biol. 2011;12:104–17.

[25] Blobel CP. ADAMs: key components in EGFR signalling and development. Nat Rev Mol Cell Biol. 2005;6:32–43.

[26] Schreier B, Gekle M, Grossmann C. Role of epidermal growth factor receptor in vascular structure and function. Curr Opin Nephrol Hypertens. 2014;23:113–21.

[27] Roberts RB, Arteaga CL, Threadgill DW. Modeling the cancer patient with genetically engineered mice: prediction of toxicity from molecule-targeted therapies. Cancer Cell. 2004;5:115–20.

[28] Nagareddy PR, Chow FL, Hao L, Wang X, Nishimura T, MacLeod KM, et al. Maintenance of adrenergic vascular tone by MMP transactivation of the EGFR requires PI3K and mitochondrial ATP synthesis. Cardiovasc Res. 2009;84:368–77.

[29] Pandey R, Shukla P, Anjum B, Gupta HP, Pal S, Arjaria N, et al. Estrogen deficiency induces memory loss via altered hippocampal HB-EGF and autophagy. J Endocrinol. 2020;244:53–70.

[30] Harraz OF, Hill-Eubanks D, Nelson MT. PIP(2): A critical regulator of vascular ion channels hiding in plain sight. Proc Natl Acad Sci U S A. 2020;117:20378–89.

[31] Fujimoto M, Shiba M, Kawakita F, Liu L, Nakasaki A, Shimojo N, et al. Epidermal growth factor-like repeats of tenascin-C-induced constriction of cerebral arteries via activation of epidermal growth factor receptors in rats. Brain Res. 2016;1642:436–44.

[32] Koide M, Penar PL, Tranmer BI, Wellman GC. Heparin-binding EGF-like growth factor mediates oxyhemoglobin-induced suppression of voltage-dependent potassium channels in rabbit cerebral artery myocytes. Am J Physiol Heart Circ Physiol. 2007;293:H1750–9.

[33] Koide M, Wellman GC. SAH-induced suppression of voltage-gated K(+) (K (V)) channel currents in parenchymal arteriolar myocytes involves activation of the HB-EGF/EGFR pathway. Acta Neurochir Suppl. 2013;115:179–84.

[34] Ishiguro M, Morielli AD, Zvarova K, Tranmer BI, Penar PL, Wellman GC. Oxyhemoglobin-induced suppression of voltage-dependent K+ channels in cerebral arteries by enhanced tyrosine kinase activity. Circ Res. 2006;99:1252–60.

[35] Hao L, Du M, Lopez-Campistrous A, Fernandez-Patron C. Agonist-induced activation of matrix metalloproteinase-7 promotes vasoconstriction through the epidermal growth factor-receptor pathway. Circ Res. 2004;94:68–76.

[36] Chansel D, Ciroldi M, Vandermeersch S, Jackson LF, Gomez AM, Henrion D, et al. Heparin binding EGF is necessary for vasospastic response to endothelin. Faseb j. 2006;20:1936–8.

[37] Wang Y, Nakayama M, Pitulescu ME, Schmidt TS, Bochenek ML, Sakakibara A, et al. Ephrin-B2 controls VEGF-induced angiogenesis and lymphangiogenesis. Nature. 2010;465:483–6.

[38] Lee TC, Threadgill DW. Generation and validation of mice carrying a conditional allele of the epidermal growth factor receptor. Genesis. 2009;47:85–92.

[39] Bethge P, Carta S, Lorenzo DA, Egolf L, Goniotaki D, Madisen L, et al. An R-CaMP1.07 reporter mouse for cell-type-specific expression of a sensitive red fluorescent calcium indicator. PLoS One. 2017;12:e0179460.

[40] Lee FK, Lee JC, Shui B, Reining S, Jibilian M, Small DM, et al. Genetically engineered mice for combinatorial cardiovascular optobiology. Elife. 2021;10.

